# On interaction of proteinoids with simulated neural networks

**DOI:** 10.1101/2023.12.01.569607

**Authors:** Panagiotis Mougkogiannis, Andrew Adamatzky

## Abstract

Proteinoids are thermal proteins which swell into microspheres in solution. The proteinoid microspheres show spiking of electrical potential similar to that to that of living neurons. Rich spectrum of proteinoids’ spiking responses to optical and electrical stimulation makes them promising candidates for neuromorphic unconventional computing devices. We decided to evaluate neuron-like activity of proteinoids in the experimental setups of their interaction with simulate neuronal network of Izhikevich neurons. The simulated neural networks stimulate and modulate electrical activity of proteinoid ensembles by interacting with them via arbitrary form programmable function generator. Different amino-acid compositions of proteinoids responded uniquely to input spiking from simulated neurons. We demonstrated that patterns of electrical spiking activity of proteinoids and complexity of the activity can be tuned by patterns of spikes generated by simulated neurons. The research opens novel venues to establishing interacting between nanobrains – brain-like organoids made from molecules, not animal cells — and real nervous systems.

## 1. Introduction

Thermal proteins, also known as proteinoids [1], are obtained by heating amino acids to their melting point and initiating polymerisation to produce polymeric chains. The polymerisation occurs at temperatures between 160–200 oC, in the absence of a solvent, initiator, or catalyst, under an inert atmosphere. Tri-functional amino acids such as glutamic acid, aspartic acid, or lysine undergo cyclisation at high temperatures and act as solvents and initiators for the polymerisation of other amino acids [2, 1]. This simple thermal condensation reaction enables the production of either acidic or basic proteinoids. When a proteinoid is immersed in an aqueous solution at moderate temperatures (approximately 50 oC), it swells and forms structures known as microspheres [1]. These microspheres are hollow and typically contain an aqueous solution. The proteinoids are capable of folding into intricate shapes and interacting with different molecules, which makes them more versatile. Proteinoids are resistant and durable, they can withstand extreme temperatures and pH levels. Proteinoids can catalyse reactions and self-assemble into larger structures.

The proteinoid microspheres exhibit a consistent membrane potential ranging from 20mV to 70mV, even in the absence of any stimulating current. Interestingly, some microspheres within the population demonstrate a steady opposite polarisation [3]. When micro-electrodes are inserted into these microspheres, electrical membrane potentials, oscillations, and action potentials can be observed. These microspheres display spikes similar to action potentials. Additionally, the electrical activity of the microspheres includes spontaneous bursts of electrical potential, referred to as flip-flops, and miniature potential activities during flopped phases [4].

While the presence of glycerol lecithin amplifies these effects, purely synthetic microspheres can still achieve a 20mV membrane potential [5]. In pure 20*μ*m microspheres, the amplitude reaches 20mV, whereas in 200*μ*m microspheres with lecithin, it can reach 70 mV. Phospholipid-free microspheres exhibit a regular amplitude of spiking [5]. Membrane, action, and oscillatory potentials have been recorded from microspheres composed of thermal protein, glycerol, and lecithin [4, 5], and these potentials have been observed for several days [6]. Furthermore, these microspheres remain stable in aqueous environments with pH levels above 7.0oC and continue oscillating for weeks [7, 3].

Neuron-like electrical activity of proteinoid microspheres makes them a promising and intriguing class of substrates to produce brain-like structures responding to a stimulation by changing patterns of their electrical spiking activity [8, 9, 10, 11, 12].

In this paper, our objective is to explore the possibility of interaction between ensembles of proteinoid microspheres and living neurons through patterns of electrical spiking activity. Answering this question could potentially pave the way for the development of sensing and computing devices utilising proteinoid microspheres that can be implanted in nervous tissue or the brain.

The collective assemblies of proteinoid microspheres hold great promise in advancing healthcare technologies and future therapeutic approaches, particularly in drug delivery and release applications [13]. This is due to their proteinlike characteristics, bio-compatibility, and non-toxicity [14]. Proteinoids are already being used to encapsulate various drugs, including methotrexate [15], hydroxyapatite [16], cholesterol [17], and drugs promoting osteogenic differentiation [18]. These diverse applications underscore the versatility and potential of proteinoid microspheres in the development of novel therapies.

If proteinoid ensembles possess the capability to respond to and interpret the electrical activity of the nervous system, it opens up possibilities for designing and prototyping intelligent drug delivery systems, further expanding their potential applications..

**Figure 1:**
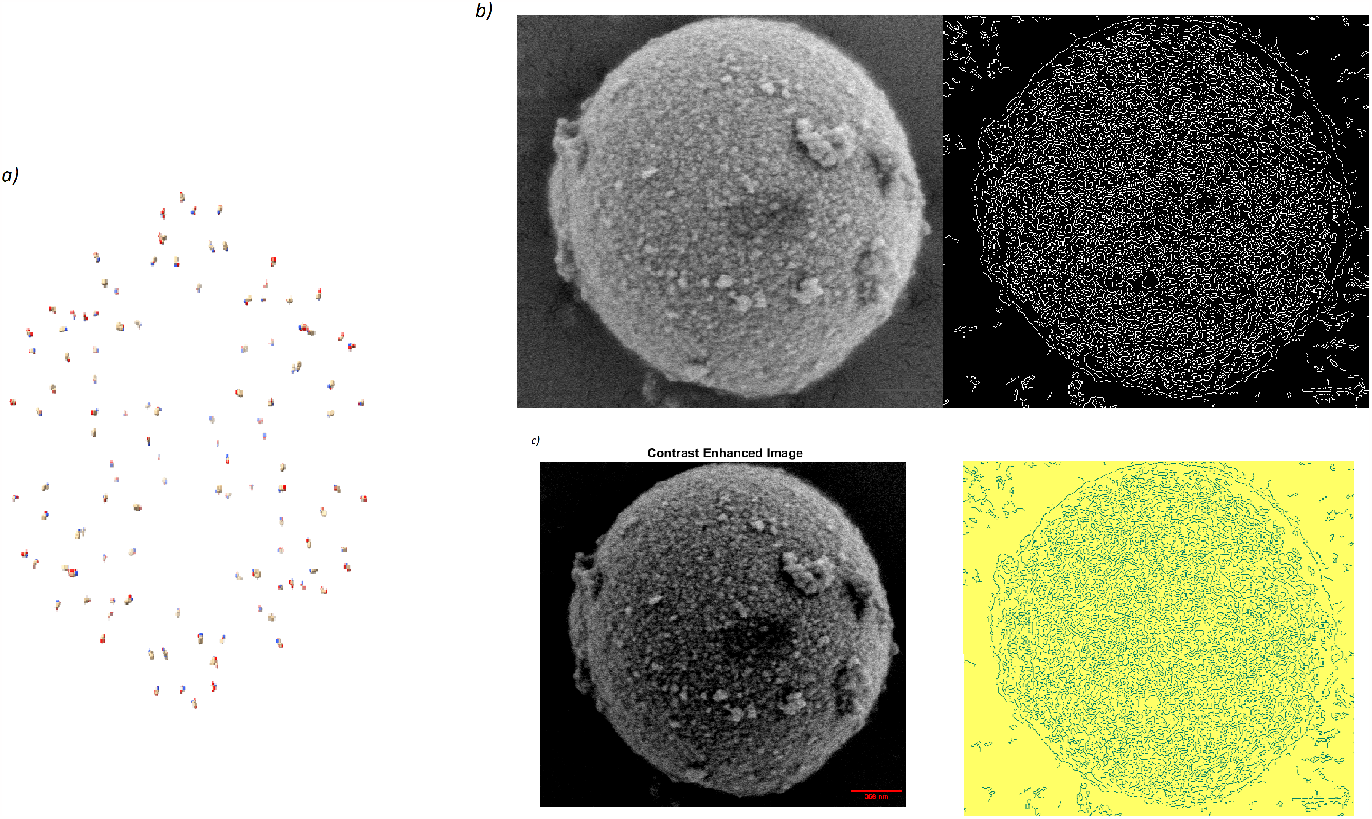
a) This image shows the spherical structure of L-Glu:L-Phe proteinoids microspher created using the AlphaFold tool and icosahedral symmetry I tools of ChimeraX. The proteinoids self-assemble into a sphere with a diameter of 1 um and the structure is stabilised by non-covalent interactions of hydrophobic and hydrophilic regions of the proteins. b) This scanning electron microscope image reveals the proteinoid nanospheres (approx. 40 nm) that comprise a vaterite microsphere with a diameter of 2257.68 nm. Applying contrast enhancement on the microscopic image exposed the edges of the microspheres. The contrast, correlation, energy, and homogeneity values were calculated as 0.2243, 0.8997, 0.1640, and 0.8879, respectively.

The paper is structured as follows. We introduce experimental techniques and computational representation of neurons in Sect. 2. We analyse co-spiking activity of ensembles of proteinoid microspheres and simulated neurons in Sect. 3. Section 4 outlines potential mechanisms on tuning patterns of electrical activity of proteinoids with spikes from simulated neurons and proposes pathways for future development.

## 2. Methods

L-Phenylalanine, L-Aspartic acid, L-Histidine, L-Glutamic acid, and L-Lysine were purchased from Sigma Aldrich and had a purity more than 98%. Proteinoids were synthesised using tried-and-true techniques indicated in [19]. Scanning electron microscopy (SEM) with FEI Quanta 650 equipment was used to examine the proteinoids’ structures. FT-IR spectroscopy was used to characterise the proteinoids [19]. Morphology of the single proteinoid microspheres is characterised in Fig. 1 and an ensemble of the proteinoid microsphers is shown in Fig. 2.

A high-resolution data logger with a 24-bit A/D converter (ADC-24, Pico Technology, UK) and iridium-coated stainless steel sub-dermal needle electrodes (Spes Medica S.r.l., Italy) were used to measure the electrical activity of the proteinoids. To assess the potential difference, pairs of electrodes were set up with a spacing of about 10 mm between them. At a rate of one sample per second, all electrical activity was captured. Multiple measurements (up to 600 per second) were captured by the data logger, and their average was saved for further study.

**Figure 2:**
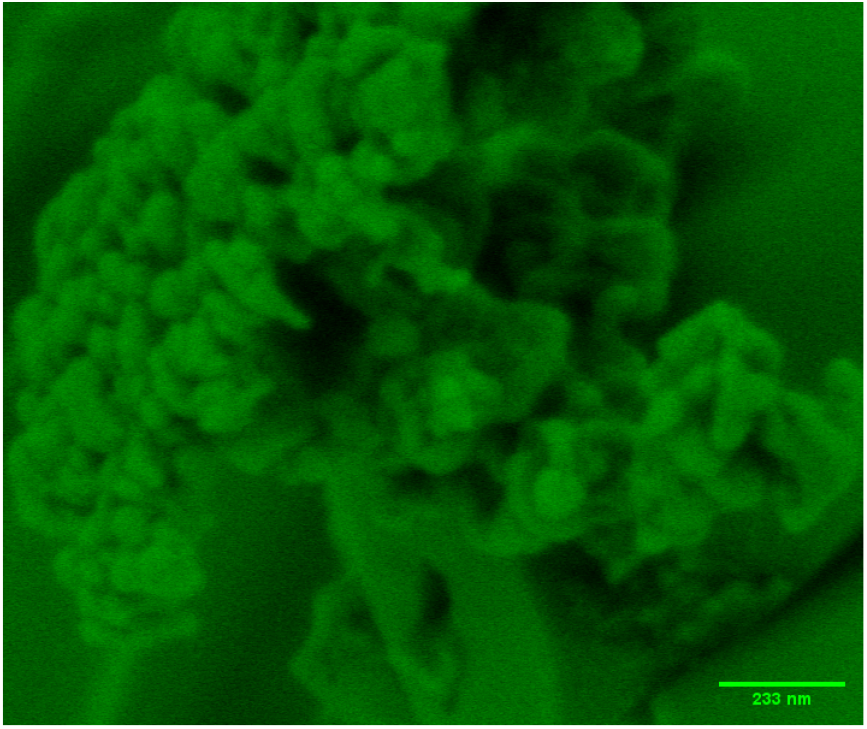
A protobrain, a self-assembled structure of proteinoids microspheres is shown in in this scanning electron microscopy (SEM) image. The ensemble spans 233 nm, as indicated by the scale bar.

We used Izhikevich model [20] to imitate neurons. The model is a biologically plausible and versatile model capable of replicating diverse spiking and bursting behaviours exhibited by actual neurons. The model comprises two ordinary differential equations that depict the membrane potential and recovery variable of a neuron. The model is made up of four parameters, namely *a, b, c*, and *d*, which regulate the structure and kinetics of the spikes and bursts (Table 1). The Izhikevich model can be written as [20]:

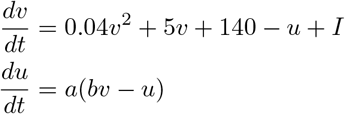

where *v* is the membrane potential, *u* is the recovery variable, *I* is the input current, and *a, b, c*, and *d* are the parameters. The model also has a reset condition that is applied whenever *v* reaches 30 mV:

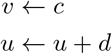

This condition emulates the sodium and potassium currents responsible for generating action potentials in actual neurons.

**Table 1:** Parameters of the Izhikevich model for different types of neurons.

| Type of neuron | a | b | c | d | I |
| --- | --- | --- | --- | --- | --- |
| Tonic spiking | 0.02 | 0.2 | -65 | 6 | 14 |
| Phasic spiking | 0.02 | 0.25 | -65 | 6 | 0.5 |
| Tonic bursting | 0.02 | 0.2 | -50 | 2 | 15 |
| Phasic bursting | 0.02 | 0.25 | -55 | 0.05 | 0.6 |
| Mixed mode | 0.02 | 0.2 | -55 | 4 | 10 |
| Spike frequency adaptation | 0.01 | 0.2 | -65 | 8 | 30 |
| Class 1 | 0.02 | -0.1 | -55 | 6 | 0 |
| Class 2 | 0.2 | 0.26 | -65 | 0 | 0 |
| Spike latency | 0.02 | 0.2 | -65 | 6 | 7 |
| Subthreshold oscillations | 0.05 | 0.26 | -60 | 0 | 0 |
| Resonator | 0.1 | 0.26 | -60 | -1 | 0 |
| Integrator | 0.02 | -0.1 | -55 | 6 | 0 |
| Rebound spike | 0.03 | 0.25 | -60 | 4 | 0 |
| Rebound burst | 0.03 | 0.25 | -52 | 0 | 0 |
| Threshold variability | 0.03 | 0.25 | -60 | 4 | 0 |
| Bistability | 1 | 1.5 | -60 | 0 | -65 |
| DAP | 1 | 0.2 | -60 | -21 | 0 |
| Accomodation | 0.02 | 1 | -55 | 4 | 0 |
| Inhibition-induced spiking | -0.02 | -1 | -60 | 8 | 80 |
| Inhibition-induced bursting | -0.026 | -1 | -45 | 0 | 80 |

This study examined the impact of a consistent current injection on the membrane potential in Izhikevich neuron. The neuron’s membrane potential and spiking frequency were recorded for each current level. Figure 3 depicts the membrane potential trace resulting from a 10 *μ*A current injection. The neuron displays tonic spiking, characterised by regular firing of action potentials.

A one-way analysis of variance (ANOVA) was conducted to determine the impact of current injection on the neuron’s spiking frequency. Table 2 presents the ANOVA results. The table displays information on the source of variation, sum of squares (SS), degrees of freedom (df), mean square (MS), F-ratio (F), and probability of obtaining a larger F-ratio by chance (Prob¿F). The study findings suggest that the spiking frequency is significantly affected by the current injection, as evidenced by the significantly larger variation between columns compared to within columns (*F* (1, 200006) = 1.0790 *×* 10^6^, *p <* 0.001).

The present injection predictably modulates the spiking frequency of the proteinoid. Increased current levels result in higher spiking frequencies.

The spiking activity recorded from Izhikevich model was fed into ensembles of proteinoid microspheres using BK 4060B function generator (B&K Precision Corporation). It is a dual channel function/arbitrary waveform generator that can generate accurate sine, square, triangle, pulse, and arbitrary waveforms, to name a few of its most prominent features and specs. It can store arbitrary waveforms in its 8 MB of memory, and it has a resolution of 16 bits. It has a maximum waveform generation rate of 75 MSa/s in real point-by-point arbitrary mode and 300 MSa/s in direct digital synthesis mode.

**Figure 3:**
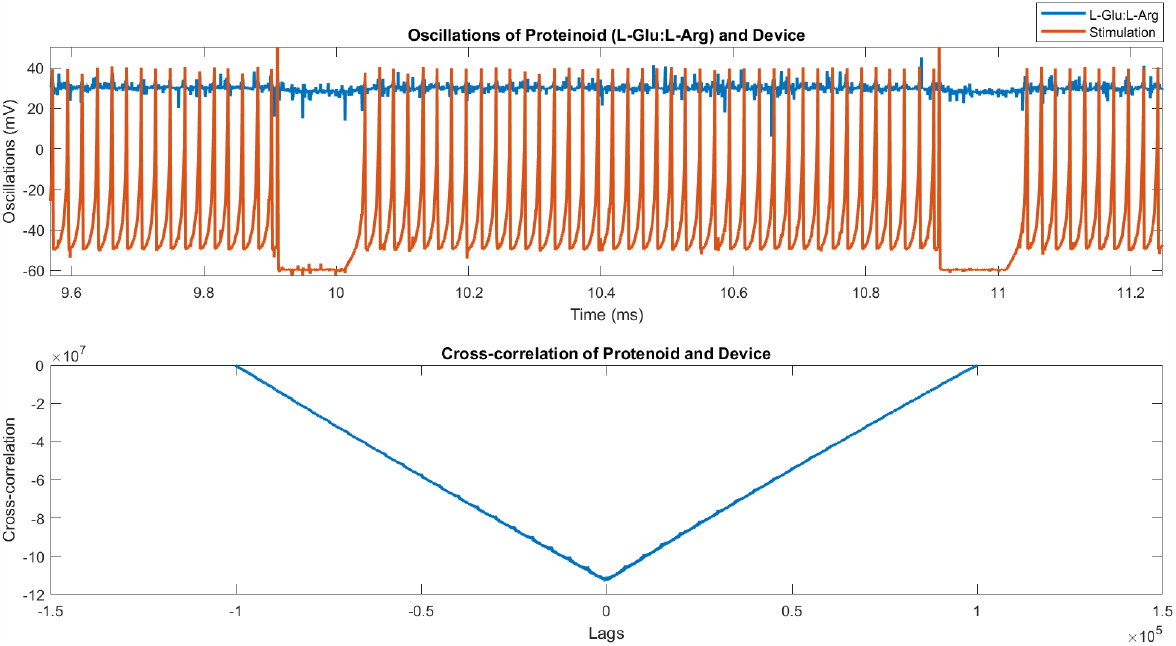
A graph depicting the membrane potential of a neuron in response to a consistent current injection. The neuron displays tonic spiking behaviour characterised by regular firing of action potentials.

**Table 2:** The table provides a statistical analysis of the tonic spiking data. The table displays information on the source of variation, sum of squares (SS), degrees of freedom (df), mean square (MS), F-ratio (F), and probability (Prob¿F) of obtaining a larger F-ratio by chance. The study shows that the difference among columns is notably greater than the difference within columns, indicating that the spiking frequency is significantly impacted by the current stimulation.

| Source | SS | df | MS | F | Prob<sub>i</sub>F |
| --- | --- | --- | --- | --- | --- |
| Columns | $2.2795 \times 10^8$ | 1 | $2.2795 \times 10^8$ | $1.0790 \times 10^6$ | 0 |
| Error | $4.2253 \times 10^7$ | 200006 | 211.2585 | - | - |
| Total | $2.7020 \times 10^8$ | 200007 | - | - | - |

## 3. Results

To examine the proteinoids L-Glu:L-Arg’s electrical characteristics and responsiveness we subjected it to stimulation with various neural spike patterns. The frequency, amplitude, peak-to-peak separation, root mean square (RMS), and standard deviation (STD) of the voltage oscillations were measured and analysed for the proteinoid and device signals. Table 3 contains a summary of the results. The findings demonstrate that the frequency and amplitude properties of the proteinoid L-Glu:L-Arg vary based on the device signal’s spike pattern, demonstrating a nonlinear and adaptive behaviour of the proteinoid material. The device frequency is between 1 Hz and 5 Hz, while the proteinoid frequency is between 3 Hz and 8 Hz. The device amplitude is between 14 mV and 90 mV, while the proteinoid amplitude is between 15 mV and 74 mV. The intricacy and diversity of the proteinoid and device signals are reflected in the voltage oscillations’ peak to peak distance, RMS, and STD, which also vary based on the spike pattern. Figure **??** illustrates the comparison of proteinoid and device frequency and amplitude for different types of spiking, as shown in Table 3.

**Figure 4:**
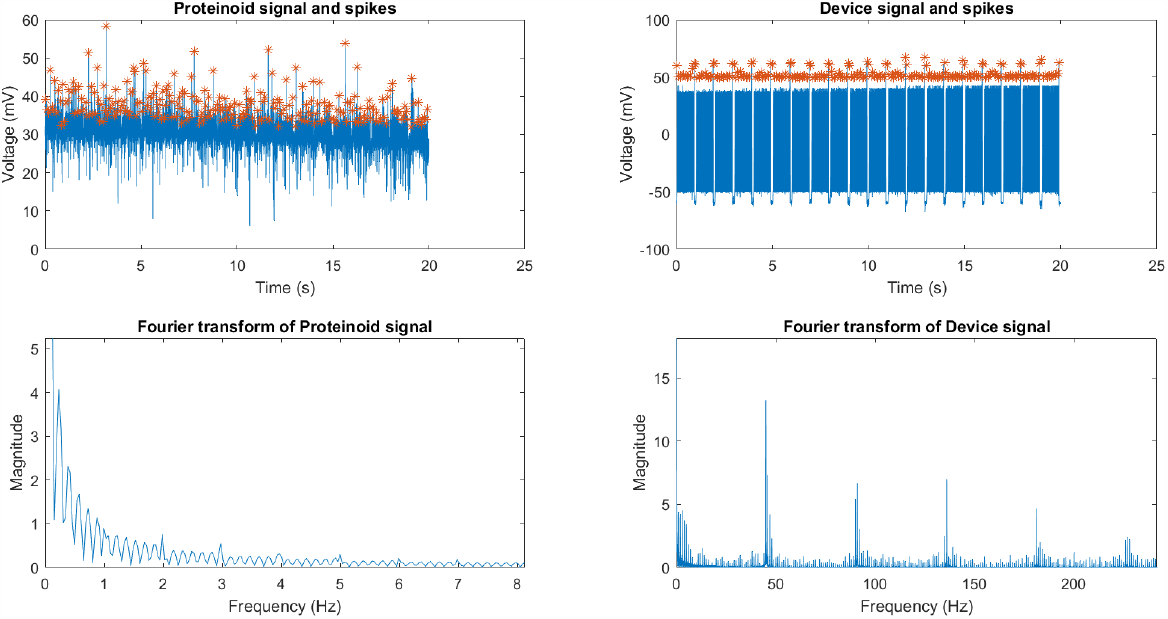
A comparative analysis of the signals and spikes produced by proteinoid and device samples. The upper panel displays the time-domain signals of proteinoid (P) and the device. Additionally, the spike trains extracted from the signals are displayed using a threshold method. The lower panel displays the frequency spectra of the signals acquired through Fourier transform. Proteinoid:L-Glu:L-Arg. Tonic spiking neurons.

**Table 3:** Frequency and amplitude of proteinoid and device spikes for different spike patterns.

| Spike Pattern | Proteinoid Frequency (Hz) | Proteinoid Amplitude (mV) | Device Frequency (Hz) | Device Amplitude (mV) |
| --- | --- | --- | --- | --- |
| Subthreshold oscillations | 7.07 | 20.81 | 1.95 | 72.90 |
| Resonator | 7.16 | 18.12 | 1.95 | 70.89 |
| Spiking Latency | 7.22 | 18.49 | 1.84 | 74.43 |
| Inhibition Induced Bursting | 7.33 | 15.57 | 3.98 | 52.18 |
| DAP | 7.06 | 21.19 | 1.33 | 77.35 |
| Thalamo Cortical | 6.85 | 14.82 | 2.84 | 70.01 |
| Integrator | 6.73 | 18.53 | 2.49 | 89.91 |
| Rebound Burst | 7.46 | 18.17 | 5.34 | 61.61 |
| Chattering | 7.04 | 18.81 | 2.07 | 79.55 |
| Mixed mode spiking | 6.46 | 14.60 | 3.88 | 70.81 |
| Tonic Spiking | 3.83 | 41.04 | 2.04 | 57.62 |
| Tonic Bursting | 7.67 | 73.59 | 3.29 | 13.92 |

Table 4 presents the measurements of peak-to-peak distance, root mean square (RMS), and standard deviation (STD) for both device and proteinoid signals across various neuronal spike conditions. The classification of neuronal spikes encompasses a total of 12 distinct types, each characterised by unique firing patterns. These patterns include subthreshold oscillations, resonator behaviour, spiking latency, and several others.

The findings indicate that there are apparent variations in the characteristics of device and proteinoid signals, contingent upon the specific type of neuronal spike under investigation. The device signal exhibits a greater peak-to-peak amplitude and lower root mean square (RMS) and standard deviation (STD) values compared to the proteinoid signal in the context of subthreshold oscillations, resonator behaviour, spiking latency, inhibition-induced bursting, depolarization after-potential (DAP), and thalamo-cortical spikes. The findings suggest that the device signal exhibits greater variability and reduced stability compared to the proteinoid signal during instances of these spikes. In contrast, it is worth noting that the proteinoid signal exhibits a greater peak to peak distance while displaying lower root mean square (RMS) and standard deviation (STD) values compared to the device signal across various spike patterns, including integrator, rebound burst, chattering, mixed mode spiking, tonic spiking, and tonic bursting spikes. The findings indicate that the proteinoid signal exhibits more fluctuation and lower stability compared to the device signal in relation to these specific spikes.

**Table 4:**
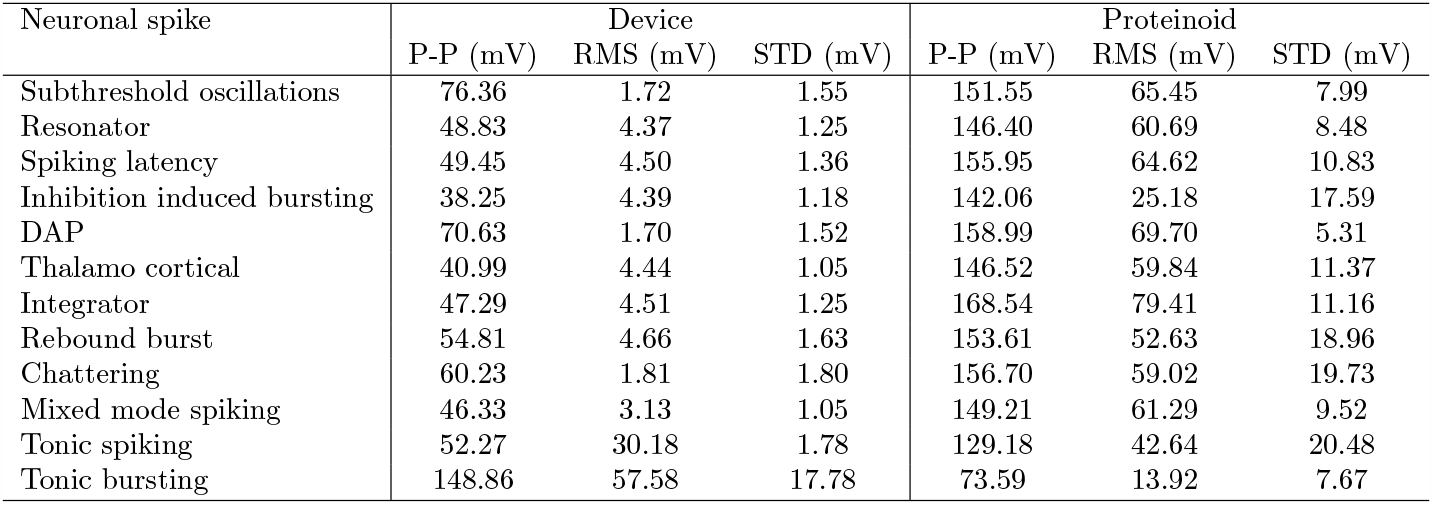
Peak to peak distance (P-P) in mV, RMS in mV, and STD of device and proteinoid signals under different neuronal spikes.

| Neuronal spike | Device |  |  | Proteinoid |  |  |
| --- | --- | --- | --- | --- | --- | --- |
|  | P-P (mV) | RMS (mV) | STD (mV) | P-P (mV) | RMS (mV) | STD (mV) |
| Subthreshold oscillations | 76.36 | 1.72 | 1.55 | 151.55 | 65.45 | 7.99 |
| Resonator | 48.83 | 4.37 | 1.25 | 146.40 | 60.69 | 8.48 |
| Spiking latency | 49.45 | 4.50 | 1.36 | 155.95 | 64.62 | 10.83 |
| Inhibition induced bursting | 38.25 | 4.39 | 1.18 | 142.06 | 25.18 | 17.59 |
| DAP | 70.63 | 1.70 | 1.52 | 158.99 | 69.70 | 5.31 |
| Thalamo cortical | 40.99 | 4.44 | 1.05 | 146.52 | 59.84 | 11.37 |
| Integrator | 47.29 | 4.51 | 1.25 | 168.54 | 79.41 | 11.16 |
| Rebound burst | 54.81 | 4.66 | 1.63 | 153.61 | 52.63 | 18.96 |
| Chattering | 60.23 | 1.81 | 1.80 | 156.70 | 59.02 | 19.73 |
| Mixed mode spiking | 46.33 | 3.13 | 1.05 | 149.21 | 61.29 | 9.52 |
| Tonic spiking | 52.27 | 30.18 | 1.78 | 129.18 | 42.64 | 20.48 |
| Tonic bursting | 148.86 | 57.58 | 17.78 | 73.59 | 13.92 | 7.67 |

A one-way analysis of variance (ANOVA) was conducted to examine the effects of different types of spiking on proteinoid L-Glu:L-Arg and the device. The dependent variables of interest included the sum of squares (SS), degrees of freedom (df), mean squares (MS), F-statistic, and p-value. The findings have been presented in Table 5. The analysis of variance (ANOVA) conducted in this study demonstrated that all types of spiking exhibited a statistically significant impact on all measures under investigation. This was evidenced by the substantial F-values observed and the correspondingly insignificant p-values obtained (*p <* 0.001 for all tests). The findings suggest that there were distinct responses observed between the proteinoid and the device when subjected to various types of spiking. Furthermore, it is noteworthy that the variability observed within each spiking type was considerably smaller compared to the variability observed between different spiking types.

As measured by S, the spike trains of proteinoid L-Glu:L-Arg and device (Figure 5) exhibited a moderate degree of synchrony over time (Figure 6). To further investigate the diversity and order of the spike trains, we computed their pairwise dissimilarities using E (Figure 6c) and arranged them using a hierarchical clustering algorithm (Figure 6d). The sorted spike trains revealed a distinct pattern of similarity among some clusters of spikes, as well as a greater degree of order than would be expected by chance (Figure 6f). The time profile E (Figure 6e) for the sorted spike trains revealed that the dissimilarity between spike trains varied over time, with periods of high and low dissimilarity. These findings indicate that the proteinoid L-Glu:L-Arg and device are capable of producing complex and dynamic spike patterns in response to tonic spiking stimuli.

**Table 5:**
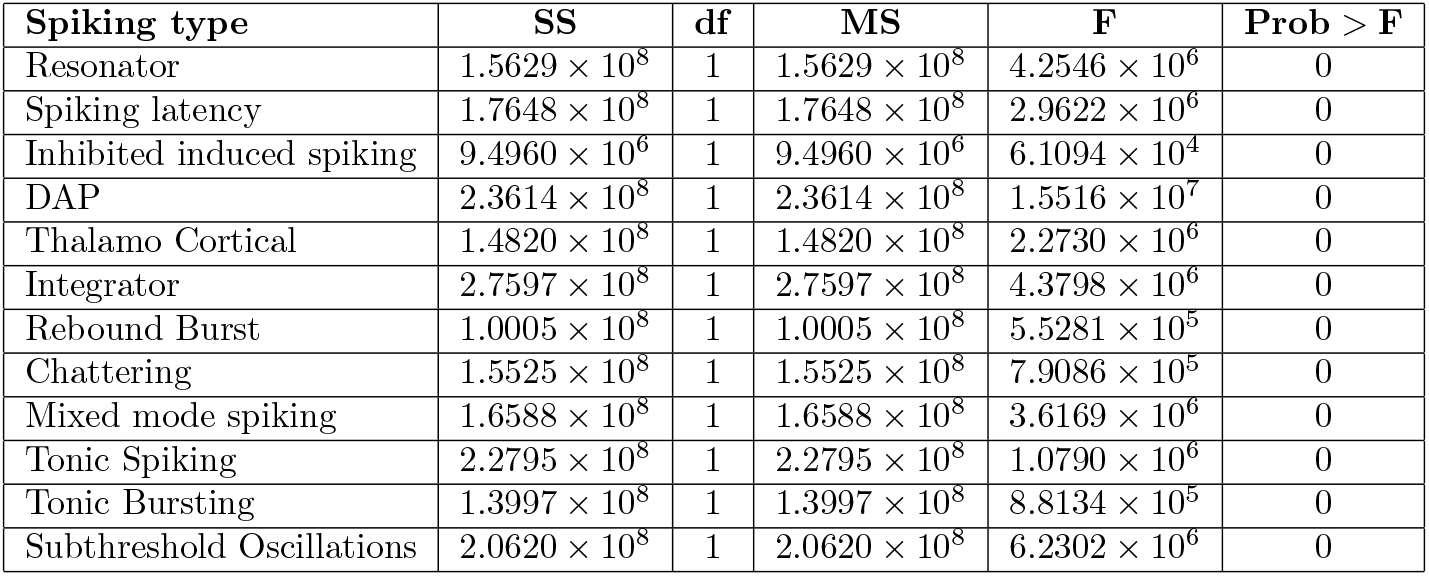
Analysis of variance (ANOVA) results for different types of spiking on proteinoid L-Glu:L-Arg and the device, showing significant differences in all measures (*p <* 0.001).

**Figure 5:**
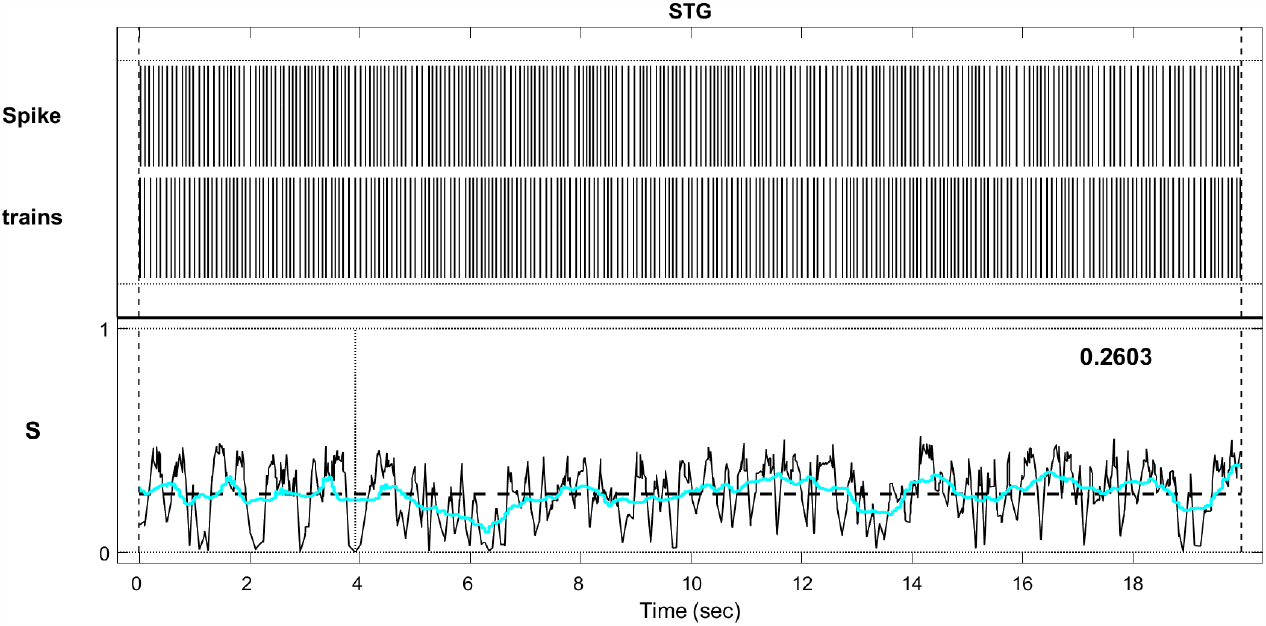
The figure depicts Spike trains of proteinoid L-Glu:L-Arg and a device utilising SPIKY matlab.The graph displays the relationship between the measure of spike train synchrony, denoted as S, and time in seconds. The mean value of S is 0.2603, suggesting a moderate degree of synchronicity among the proteinoid spikes.

The spike pattern’s effect on the Lempel-Ziv complexity (LZC) of the resulting binary sequence was one of the primary research questions in this work. By determining the shortest description of the sequence by compression using repeating patterns, the LZC provides a measure of the sequence’s randomness or unpredictability. A greater LZC indicates that compressing the sequence will be more of a challenge. Figure 19, depicts the normalised LZC of the binary sequence of spikes for various neural dynamics. To provide uniformity in comparisons across sequence lengths, the LZC was divided by the sequence length. Different forms of spike patterns exhibit varying degrees of complexity and unpredictability in the LZC, as seen in the Figure 19. LZC values range from 0.95 for tonic bursting to 0.15 for subthreshold oscillations. This indicates that tonic bursting has more unpredictability and diversity in its spiking behaviour than subthreshold oscillations, making it more difficult to compress and characterise. Subthreshold oscillations, on the other hand, are more stable and regular, with a more straightforward description. LZC values range from 0.35 to 0.65 for the other spike pattern types. These findings suggest that the spike pattern can serve as an indicator for characterising the complexity of neural dynamics, with the LZC serving as a measure for doing so [21, 22, 23, 24].

**Figure 6:**
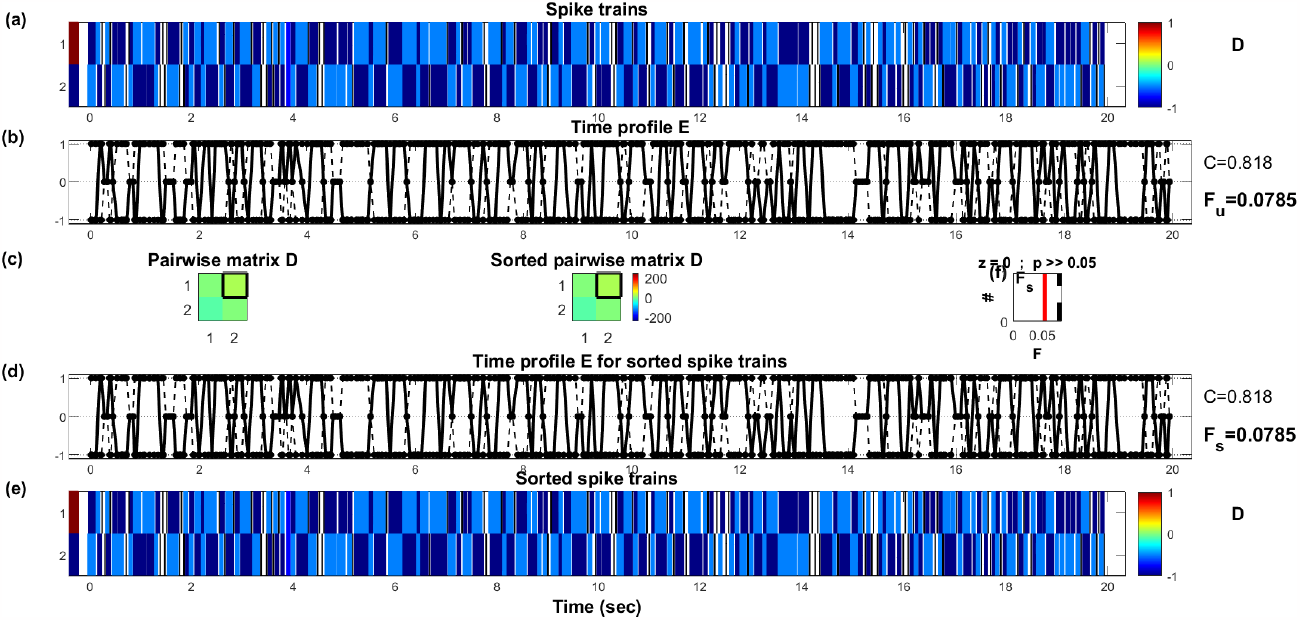
a) Spike trains of L-Glu:L-Arg proteinoid and device using SPIKY matlab. b) Time profile E with C equal to 0.81 and Fu equal to 0.0785. The graph depicts the relationship between E, a measure of spike train dissimilarity, and time in seconds. C is the average dissimilarity over the entire time interval, and Fu is the fraction of the time when E is greater than a predetermined threshold. E quantifies the dissimilarity between two spike trains at each instant, with higher values indicating greater dissimilarity. c) Pairwise matrix D. The matrix displays the pairwise dissimilarities between every spike train in the dataset, as computed by E. The dissimilarities are color-coded from blue (low) to red (high) to indicate their degree. The matrix is diagonally symmetrical. d) Pairwise-sorted matrix D. The matrix displays the same pairwise dissimilarities as shown in c), but is arranged according to a hierarchical clustering algorithm that clusters spike trains with similar characteristics. The top dendrogram displays the clustering structure and distances between clusters. f) z=0 p¿¿0.05 graph F. The graph depicts the relationship between F, a measure of spike train order, and time in seconds. F measures how ordered or chaotic a collection of spike trains is at each instant in time, with greater values indicating greater order. The horizontal line at z=0 represents the level of order expected by chance, while the shaded region represents the p-value for testing the null hypothesis that F equals z. e) Time profile E for ordered spike trains with C = 0.81 and Fs = 0.0785. The graph depicts the same value of E as in (b), but for the spike trains sorted according to (d). Fs represents the fraction of time that E is above a predetermined threshold for the sorted spike trains.

**Figure 7:**
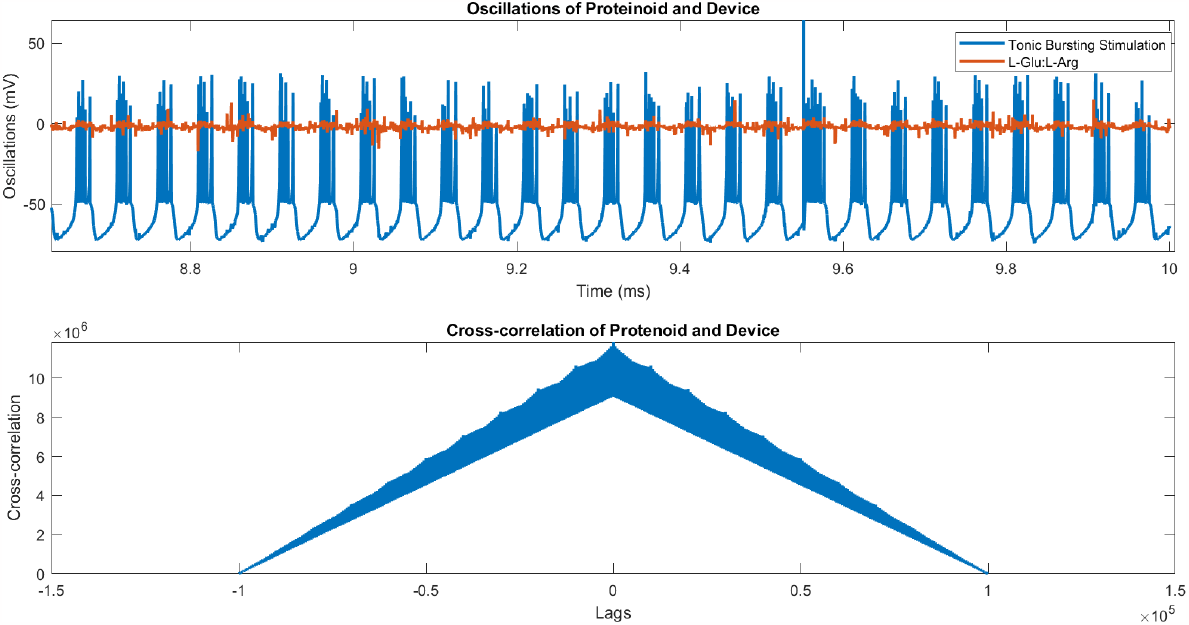
This figure shows the effects of tonic bursting stimulation on the cross-correlation of oscillation between L-Glu:L-Arg proteinoids and a device. The applied stimulation pattern is characterised by a fixed burst duration of 1000 ms and a variable inter-burst interval, exhibiting a tonic bursting mode. The figure displays the lags plot of the cross-correlation between proteinoid and device spikes at different stimulation frequencies. The lags plot displays the optimal lag value, measured in milliseconds, that maximises the cross-correlation coefficient for every column. The lags plot demonstrates that the proteinoids exhibit a variable delay in response to device spikes, which is dependent on the frequency of stimulation.

**Table 6:** A comparison of the parameters of the spike train for L-Glu:L-Arg proteinoids during various stimulation bodes. The spike train parameters C (coefficient of variation), Fu (fraction of uncorrelated spikes), and S (synchrony index) are calculated for L-Glu:L-Arg proteinoids in response to five distinct stimulation modes: tonic bursting, tonic spiking, mixed mode spiking, chattering spiking, rebound burst, integrator, thalamo cortical, dap, inhibition induced spiking, spike latency, resonator, and subthreshold oscillations.

| Stimulation mode | C | Fu | S |
| --- | --- | --- | --- |
| Tonic bursting | 0.570 | 0.0388 | 0.2980 |
| Tonic spiking | 0.818 | 0.0785 | 0.2603 |
| Mixed mode spiking | 0.594 | 0 | 0.2858 |
| Chattering spiking | 0.690 | 0.0423 | 0.2512 |
| Rebound Burst | 0.731 | 0.0897 | 0.2525 |
| Integrator | 0.747 | -0.0232 | 0.2153 |
| Thalamo Cortical | 0.625 | 0 | 0.2610 |
| DAP | 0.272 | -0.0988 | 0.3464 |
| Inhibition Induced Bursting | 0.630 | -0.1110 | 0.2538 |
| Spike Latency | 0.400 | -0.0842 | 0.2929 |
| Resonator | 0.341 | 0.0488 | 0.3579 |
| Subthreshold Oscillations | 0.414 | -0.023 | 0.3184 |

**Figure 8:**
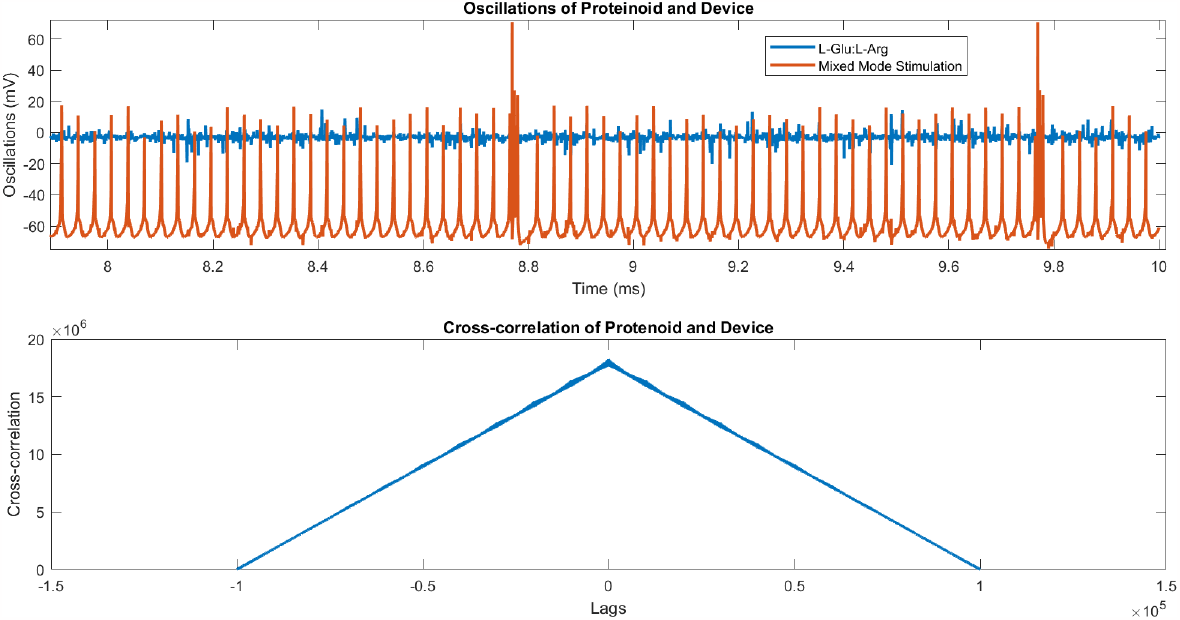
This figure shows the oscillations of L-Glu:L-Arg proteinoids and their correlation with mixed mode neuron spiking and cross-correlation lag. The voltage traces of the proteinoids and the device are depicted in the figure for varying stimulation frequencies. The stimulation pattern comprises a variable burst duration and inter-burst interval in a mixed mode spiking mode. The figure displays the cross-correlation lags plot between proteinoid and device spikes for various stimulation frequencies. The lag plot displays the optimal lag value, measured in milliseconds, that maximises the cross-correlation coefficient for every column. The figure displays the statistical results of testing the null hypothesis of no linear relationship between proteinoids and the device. The cross-correlation coefficient (R) is 0.4670, the hypothesis test result (h) is 1, and the p-value (p) is 0. The figure illustrates that the proteinoids have a mean voltage of -2.9476 mV, whereas the device has a mean voltage of -60.5448 mV. The figure illustrates a moderate positive linear correlation between proteinoids and the device. Additionally, the proteinoids exhibit a delayed response to device spikes that varies according to the frequency of stimulation.

**Figure 9:**
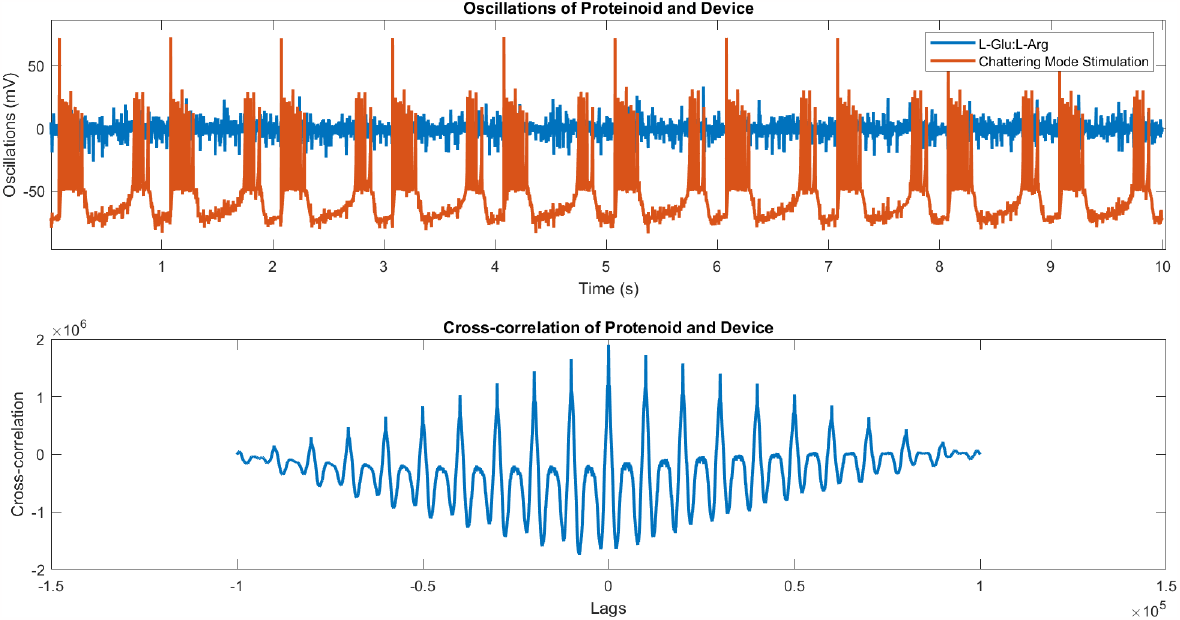
The potential of L-Glu:L-Arg and the device in Chattering spiking mode, as well as the correlation between the two. (A) The L-Glu:L-Arg (blue) and device (red) potentials in Chattering spiking mode. The respective mean values are 0.0976 mV and -55.6233 mV. (B) The cross correlation graph of L-Glu:L-Arg and the device’s potential. At latency 0, the maximum value is 0.6862, indicating a strong positive correlation. The h=1 and p=0 test results indicate that the correlation is statistically significant.

**Figure 10:**
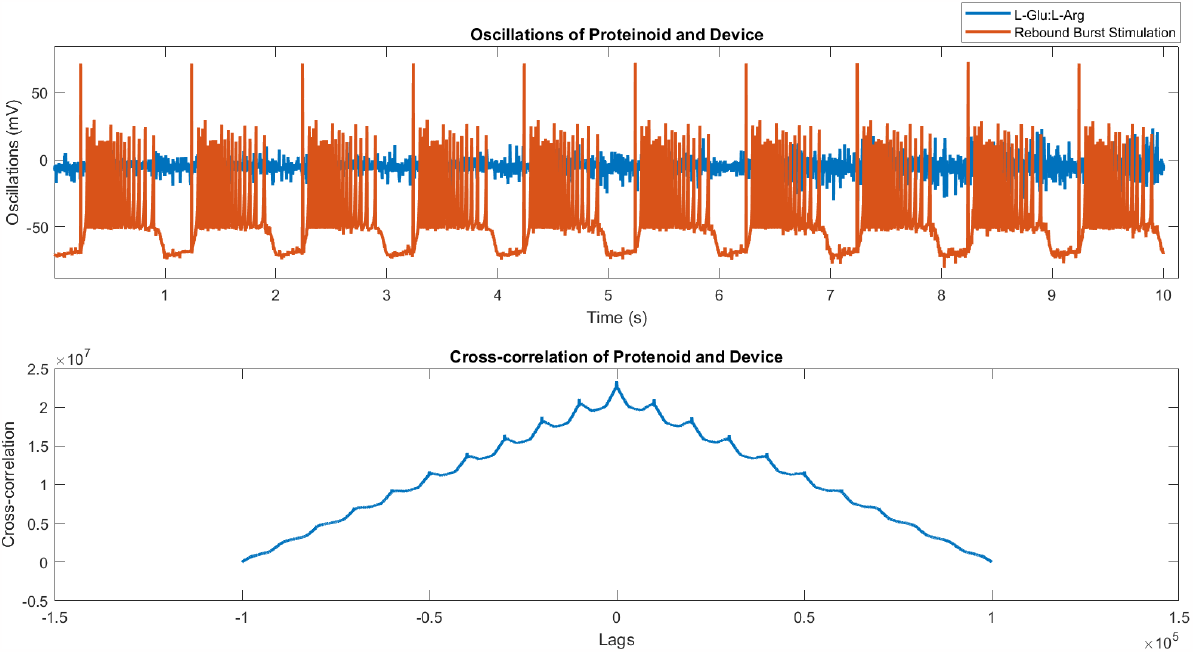
a) The potential of L-Glu:L-Arg and a device operating under Rebound Burst mode, as well as the cross correlation of their signals, are being investigated. b) The L-Glu:L-Arg compound and device exhibit periodic oscillation with a peak-to-peak amplitude of approximately 50 mV when operating under Rebound Burst mode. The potential signals of L-Glu:L-Arg and the device exhibit a positive correlation with a maximum value of 0.6506 at zero lag, as indicated by their cross correlation function. The dashed horizontal line denotes the significance level of p=0.05. The mean potential signals for L-Glu:L-Arg and the device are -4.3644 mV and -49.0963 mV, respectively.

**Figure 11:**
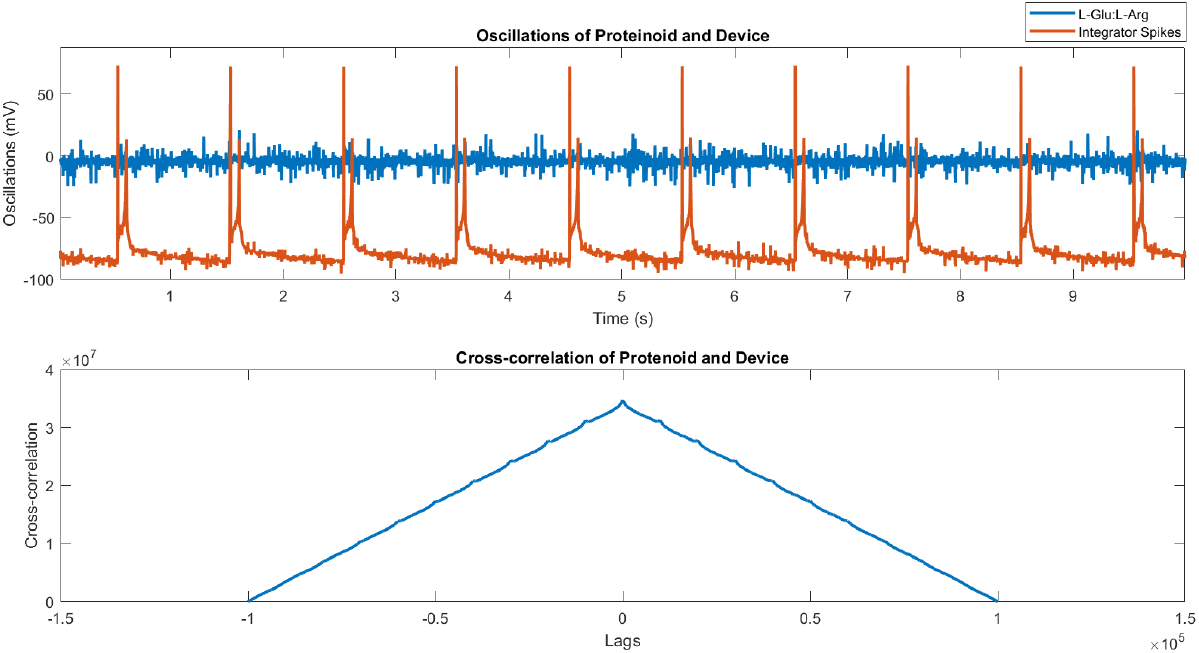
This figure shows the oscillations of proteinoids (L-Glu:L-Arg) and a device under integrator spike behaviour. The blue line represents the potential of proteinoids (L-Glu:L-Arg), which are abiotically formed protein-like molecules derived from amino acids. The red line depicts the device’s potential. The cross correlation of the two potentials in the bottom panel indicates a significant level of synchrony (R=0.5430, P¡0.001). The proteinoids had a mean voltage of -4.3285 mV, while the device had a mean voltage of -78.6202 mV.

**Figure 12:**
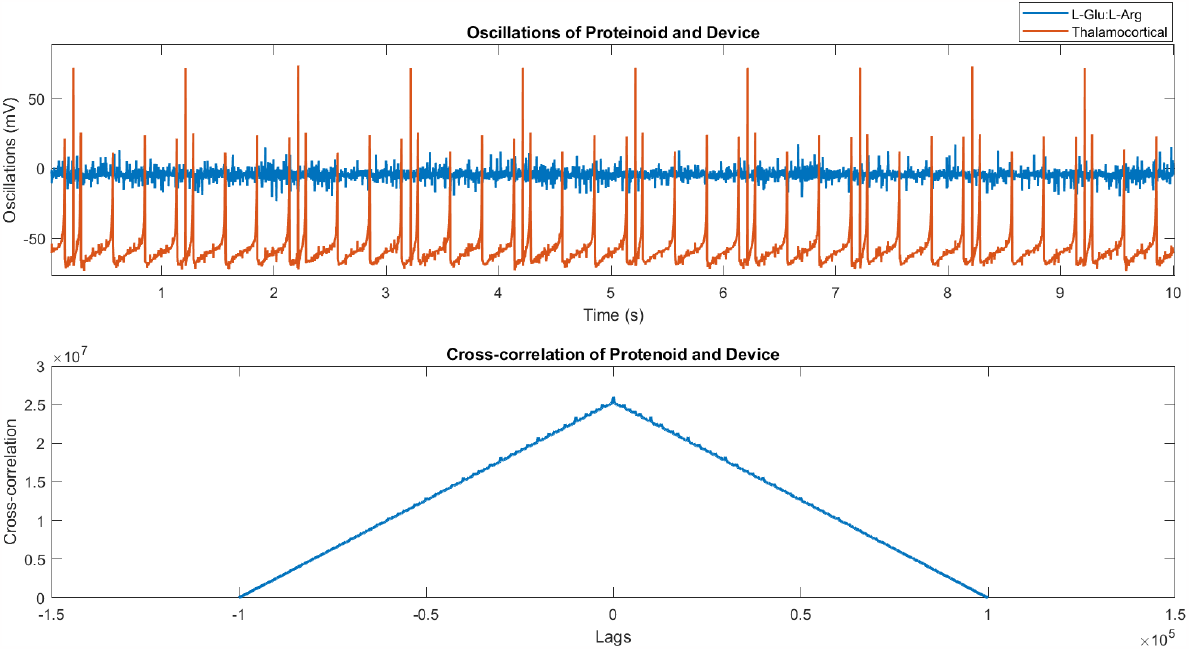
Proteinoid L-Glu:L-Arg spiking in the thalamic-cortical spiking and the device. Time-varying voltage readings from the proteinoid and the device are depicted in the graphs. When compared to the proteinoid’s regular spiking pattern and mean voltage of -4.3115 mV, the device’s spiking pattern and mean voltage of -58.7536 mV are more erratic and noisy. The proteinoid and the device have a somewhat favourable linear relationship, as indicated by the correlation coefficient of 0.5754.

**Figure 13:**
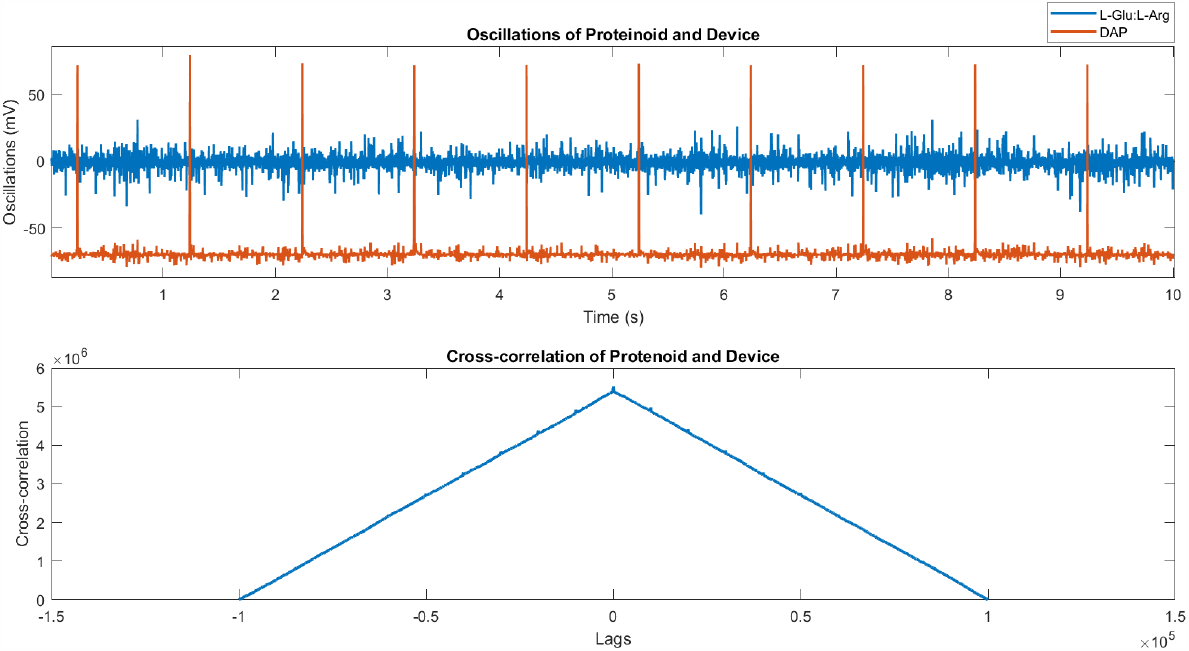
Oscillations of L-Glu:L-Arg and the device in response to DAP stimulation. The top panel displays the voltage time series of L-Glu:L-Arg (blue) and the device (red), with respective mean values of -0.7768 mV and -69.4975 mV. The cross-correlation function between the two signals has a maximal value of 0.1543 at lag zero, indicating a weakly positive correlation. The correlation is statistically significant based on the test result h=1 and the p-value of 0.

**Figure 14:**
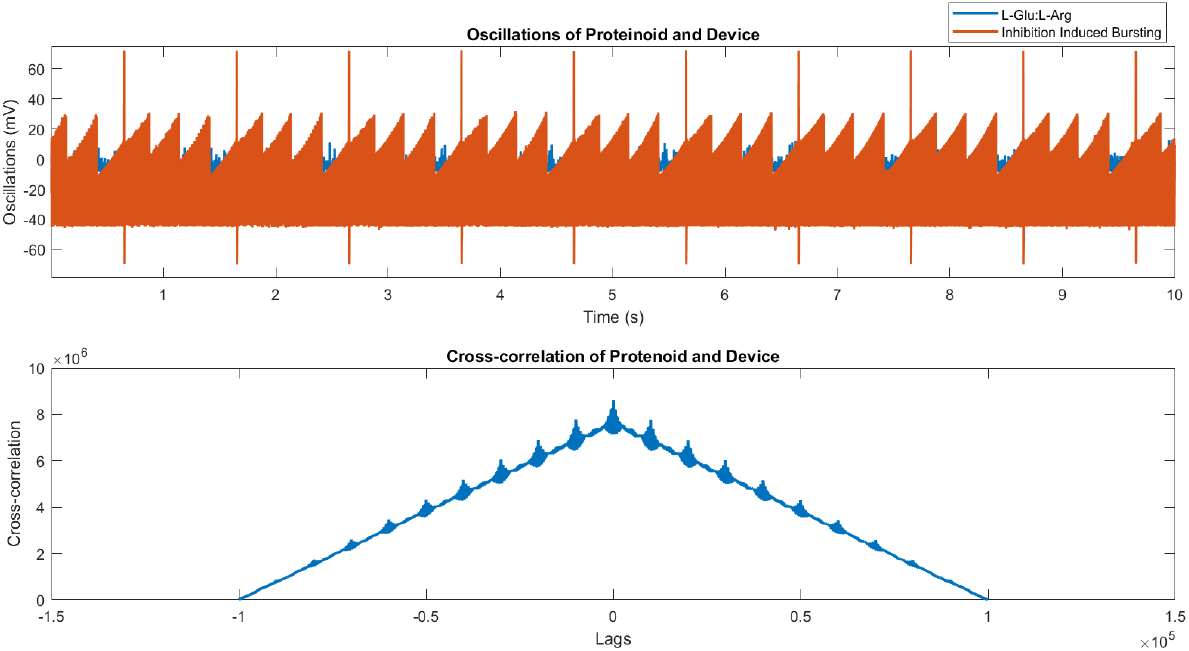
The response of L-Glu:L-Arg and the device to inhibition-induced bursting, as well as the cross-correlation between their mV potential values. The correlation was statistically significant, with R=0.4846, indicating a moderately positive relationship (h=1, p=0). The proteinoid had a mean potential of -4.2289 mV, while the device had a mean potential of -18.0097 mV.

**Figure 15:**
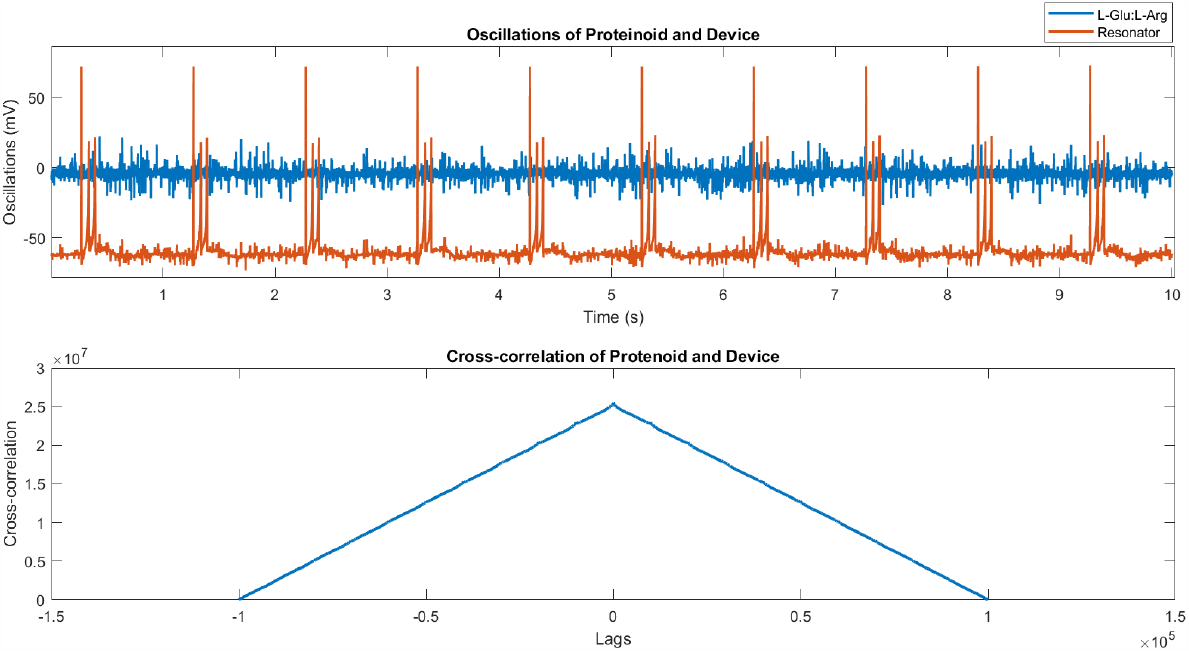
The response of L-Glu:L-Arg and the device to resonator spiking mode, as well as the cross-correlation between their mV potential values. The correlation was statistically significant, with R=0.3608, indicating a moderately positive relationship (h=1, p=0). The proteinoid had a mean potential of -4.1880 mV, while the device had a mean potential of -60.0954 mV.

**Figure 16:**
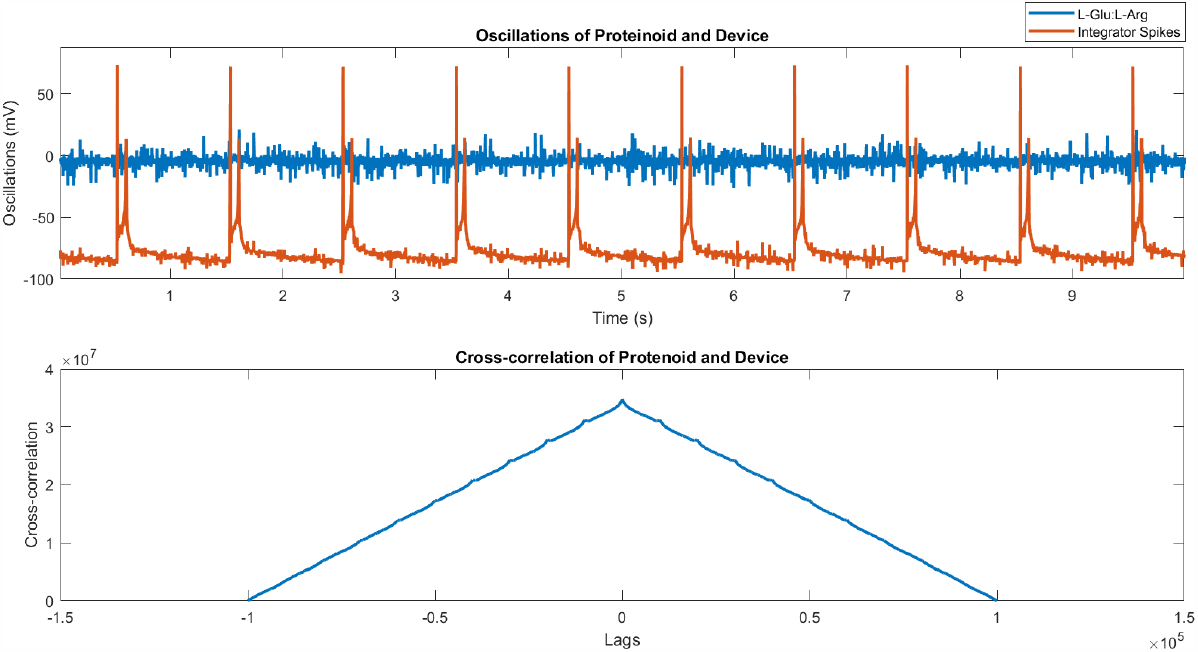
The presented figure illustrates the oscillatory behaviour of proteinoids composed of L-Glu:L-Arg, as well as a device exhibiting subthreshold oscillations. The blue line in the graph illustrates the potential of proteinoids (specifically, L-Glu:L-Arg), which are proteinlike molecules that are formed abiotically from amino acids. The red line serves as a visual representation of the potential exhibited by the device. The analysis of the cross correlation between the two potentials displayed in the lower panel reveals a statistically significant positive correlation (R=0.2658, P¡0.001). The proteinoids exhibited an average voltage of -0.7456 millivolts, whereas the device demonstrated an average voltage of -64.9624 millivolts.

**Figure 17:**
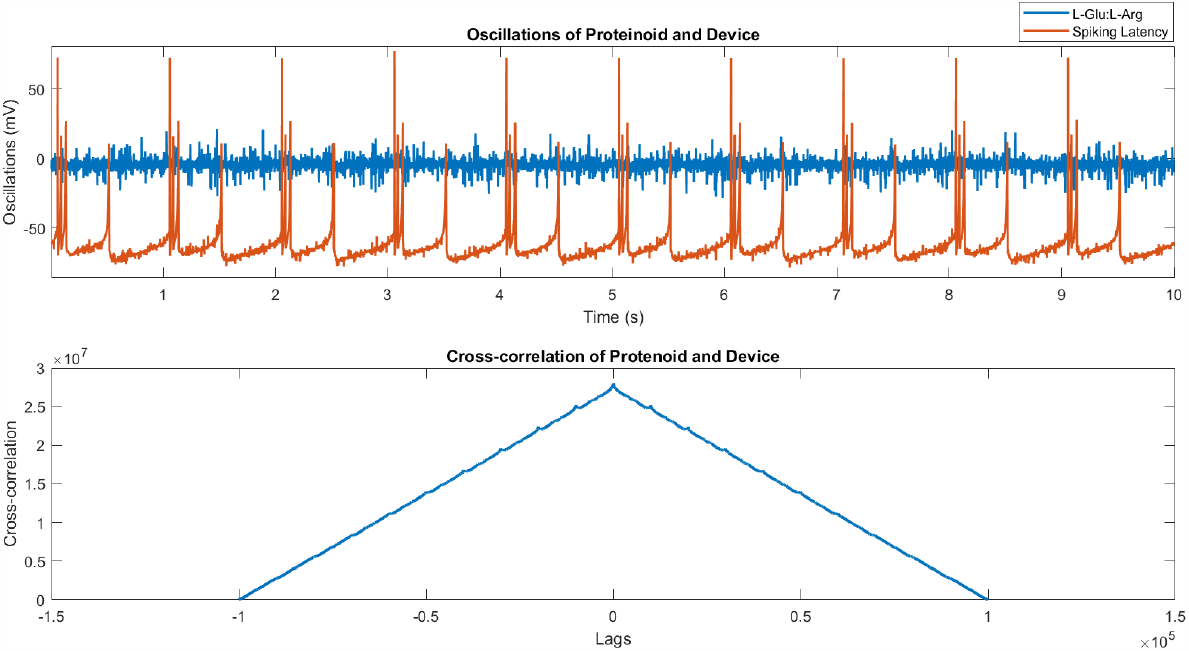
The responses of L-Glu:L-Arg and a device to spike latency neuronal stimulation showed a positive correlation between their potentials (R=0.4441), with a mean voltage of proteinoid = -4.2945 mV and a mean voltage of device = -63.7044 mV, respectively, and a p-value of 0.

**Figure 18:**
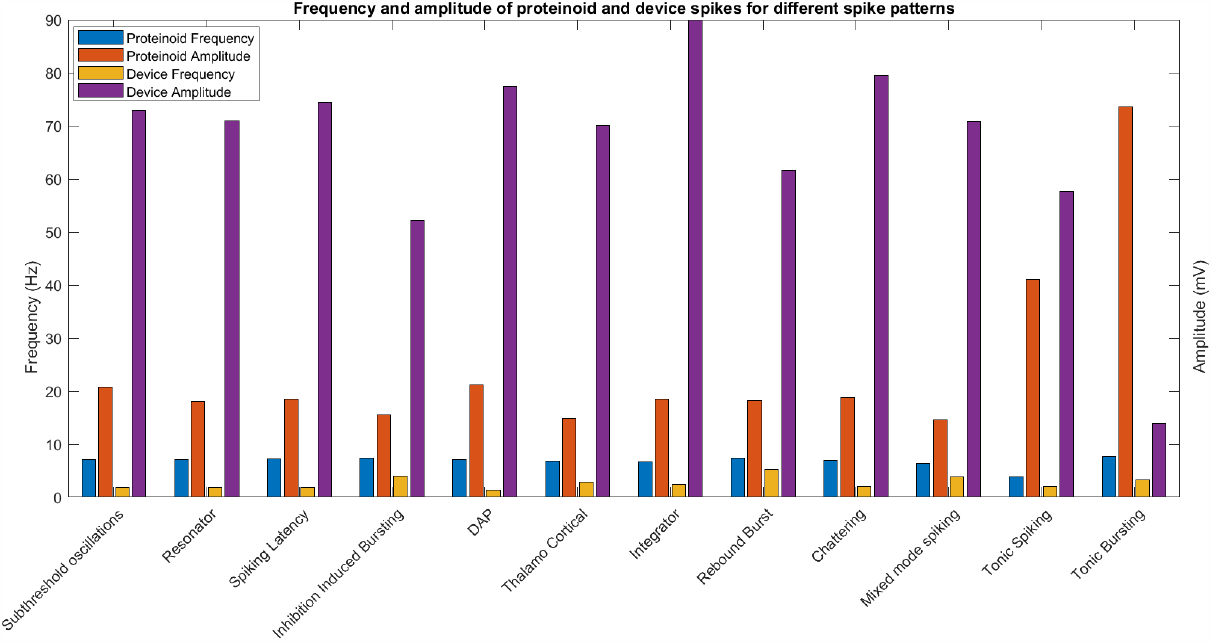
Comparison of the frequency and amplitude of various types of proteinoid and device spiking. The bar chart displays the mean frequency (Hz) and amplitude (mV) values for each spiking type, as measured from PicoScope 4000 series using BK Precision 4053 B transfer voltage generator. Proteinoid frequency is greatest for tonic bursting (7.67 Hz) and lowest for tonic spiking (3.83 Hz), whereas proteinoid amplitude is greatest for tonic bursting (73.59 mV) and lowest for thalamo cortical (14.82 mV). The device frequency is greatest during rebound burst (5.34 Hz) and lowest during DAP (1.33 Hz), whereas the device amplitude is highest during integrator (89.91 mV) and lowest during tonic bursting (13.1 mV). The results indicate that the proteinoid and device respond differently to various forms of spiking, with some types producing more pronounced effects than others.

Figure 20 displays the normalised LZC (Lempel-Ziv complexity) of the binary sequences obtained from the proteinoid frequency, proteinoid amplitude, device frequency, and device amplitude columns in the data matrix, plotted against their respective variables. Normalisation was achieved by dividing the LZC (Lempel-Ziv Complexity) by the length of the sequence, thereby enabling comparability across sequences of varying lengths. The figure demonstrates that there is no significant correlation between the LZC and the variables, except for a weak negative correlation between the LZC and the device frequency (R = -0.34). This suggests that the complexity of the binary sequences is primarily determined by their patterns and variations, rather than being strongly influenced by the values of the variables. The figure demonstrates that various spike patterns exhibit distinct distributions of LZC and variables, enabling their differentiation. Tonic bursting exhibits low LZC and device frequency, but high proteinoid frequency and amplitude. In contrast, subthreshold oscillations display high LZC and device frequency, but low proteinoid frequency and amplitude.

**Figure 19:**
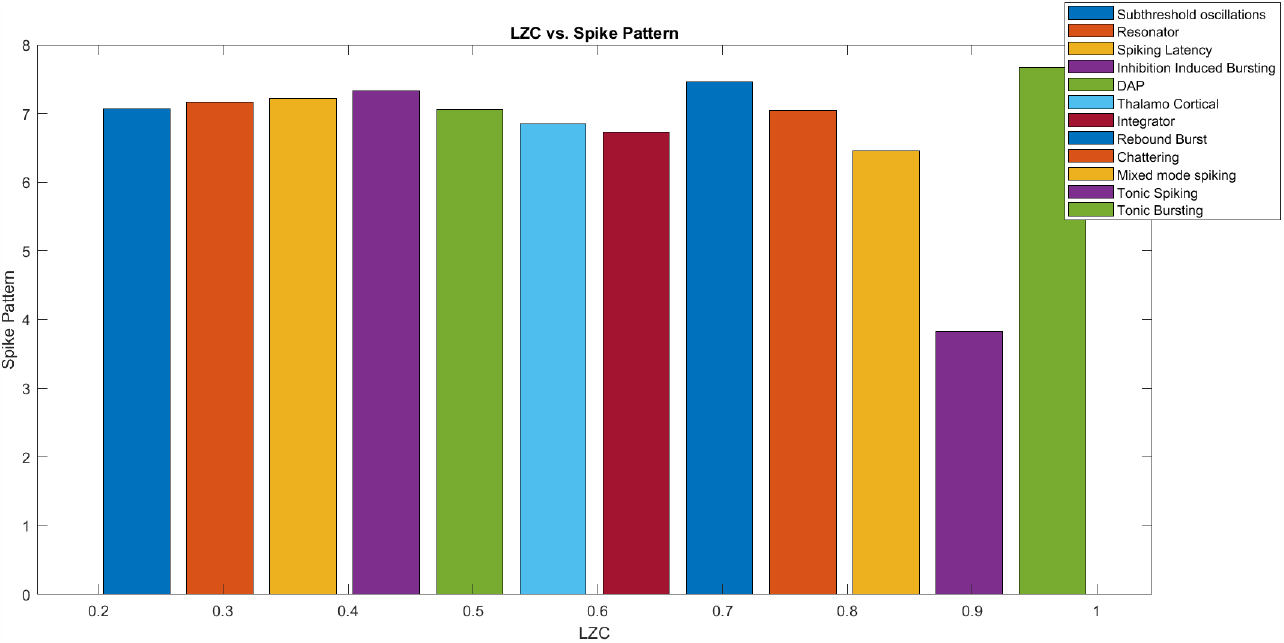
Different types of neural dynamics are represented in the bar chart by their respective normalised Lempel-Ziv complexity (LZC) of the spike pattern binary sequence. Using a threshold of 10 in the first column of the data matrix yields the spike pattern. The exhaustive approach is used to compute the LZC, and then it is normalised by dividing the result by the sequence length. Each sort of spike design is identified by name in the legend. differed types of spike patterns have varied LZCs, as seen in the above image, with more complicated and unexpected patterns having higher values. The LZC is highest for tonic bursts and lowest for subthreshold oscillations. Compared to subthreshold oscillations, this shows that tonic bursting is more difficult to compress and characterise.

**Figure 20:**
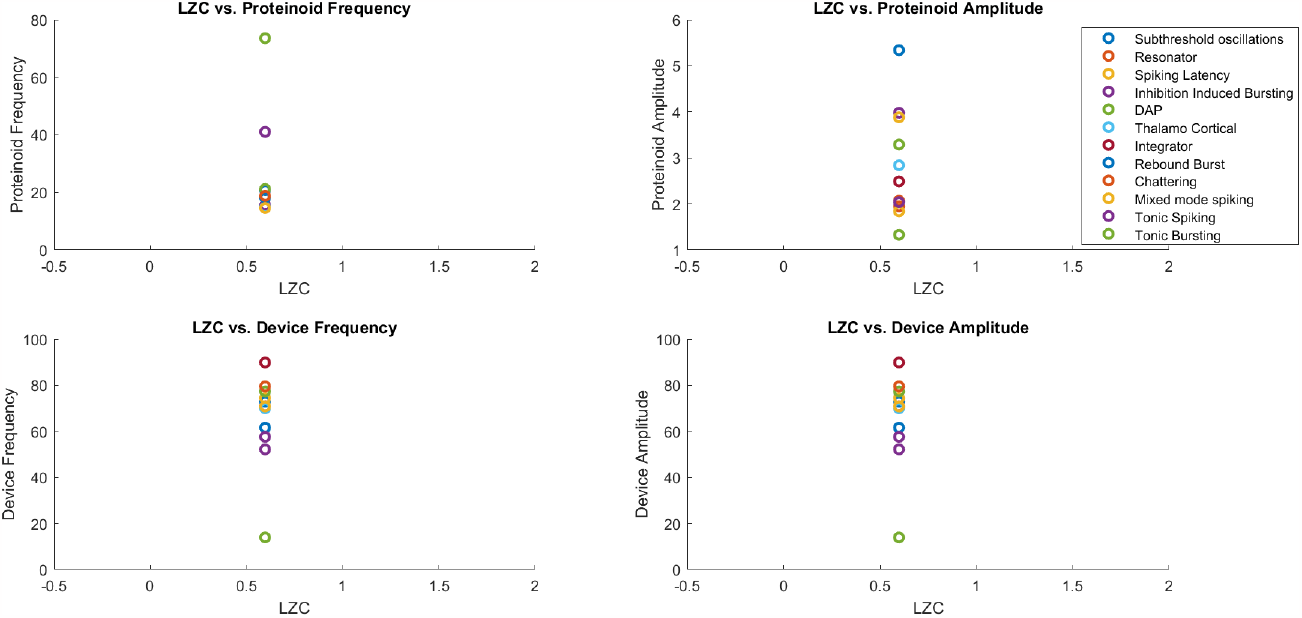
The scatter diagrams depict the normalised Lempel-Ziv complexity (LZC) of binary sequences derived from the proteinoid frequency, proteinoid amplitude, device frequency, and device amplitude columns of the data matrix against the corresponding variables. The LZC is computed using the exhaustive method and normalised by dividing it by the sequence’s length. The legend identifies each variety of spike pattern. There is no distinct correlation between the LZC and the variables, with the exception of a weak negative correlation between the LZC and device frequency. This suggests that the complexity of binary sequences is influenced less by the values of the variables and more by their patterns and variations. Different types of spike patterns have distinct distributions of LZC and variables, which can be used to differentiate them from one another. In contrast, subthreshold oscillations have high values of LZC and device frequency, but low values of proteinoid frequency and amplitude.

**Figure 21:**
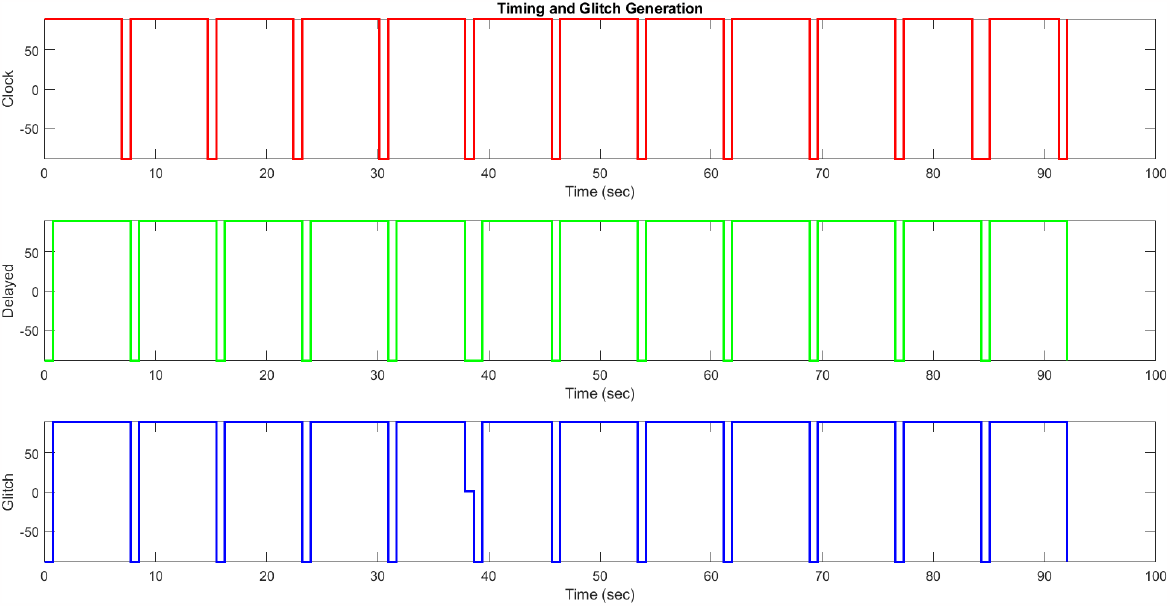
The scatter plots depict the timing and glitch generation of the proteinoid frequency, device frequency, and device amplitude signals. The proteinoid frequency serves as the clock signal, the device frequency acts as the delayed signal, and the device amplitude functions as the glitch signal. The clock signal has a frequency of 7.67 Hz and a duty cycle of 0.51. The delayed signal is a version of the clock signal that has been shifted by a delay of 2.76 Hz. The glitch signal is formed by the addition of a delayed signal and a pulse, where the pulse has a width of 0.14. The figure illustrates that the glitch arises during the transition of the delayed signal from a low to high state. Additionally, it demonstrates that the amplitude of the glitch varies depending on the amplitude value of the device.

The mean LZC value for all binary sequences was 0.597494, suggesting a moderate degree of complexity and randomness. The LZC’s standard deviation was 0.215526, indicating significant complexity variation among spike patterns. Tonic spiking exhibited the highest LZC value (0.95), indicating its complex and unpredictable nature, as well as its diverse and variable spiking behaviour. The subthreshold oscillations exhibited the lowest LZC value (0.15), indicating their simplicity and predictability, allowing for a straightforward pattern description. The results suggest that the LZC is a useful feature for characterising neuronal dynamics complexity. Additionally, it can capture certain aspects of spiking behaviour that are not apparent from the variables alone.

In addition, the timing and glitch generation of proteinoid frequency, device frequency, and device amplitude signals were investigated. Utilising a square wave as a clock signal, a shifted version of the clock signal as a delayed signal, and a combination of the delayed signal and a pulse as a glitch signal formed the basis of the timing and glitch generation. Figure 21 depicts scatter plots of these signals for various spike pattern categories. Maximum and mean values of the proteinoid frequency determined the period and duty cycle of the clock signal, which were 7.67 Hz and 0.51 respectively. The average frequency of the device determined the delay of 2.76 Hz for the delayed signal. The extent of the glitch signal was determined by the standard deviation of the device’s amplitude, which was 0.14.

The diagram reveals that the timing and generation of glitches varied across spike pattern types, with some having more frequent and larger malfunctions. For instance, tonic bursting consisted of a single glitch with a small amplitude, whereas chattering contained four anomalies with large amplitudes. This suggests that timing and glitch generation can be used to characterise the spiking behaviour of various types of neuronal dynamics, and that they can reflect aspects of the proteinoid frequency, device frequency, and device amplitude signals that cannot be captured by their values alone [25].

**Figure 22:**
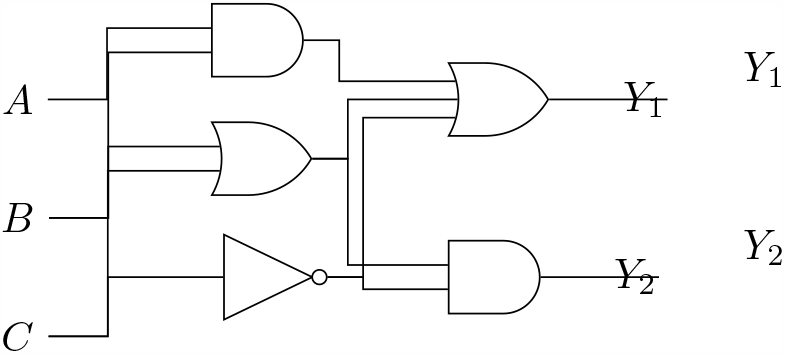
*Y*_1_ and *Y*_2_ represent the two outputs of a three-level logic system depicted in this diagram. The system has three inputs, *A, B*, and *C*, as well as six logic gates, including two AND gates, two OR gates, and two NOT gates.The diagram illustrates the connections between inputs, gates, and outputs.

In this study, six logic gates were used to design and implement a threelevel logic system with two outputs, *Y*_1_ and *Y*_2_. Figure 22 displays the system’s schematic diagram. Three inputs, *A, B*, and *C*, can be either 0 or 1: *A, B*, and *C*. These logical expressions determine the outputs:

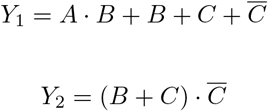

We evaluated the system with various input combinations and confirmed that the outputs corresponded to the expected values. Table 22 presents a summary of the results.

The primary objective of this investigation was to examine the spiking patterns exhibited by various neuronal types and explore the potential correlation between device potential and proteinoids. A Izhikevich spiking neuron model was employed in this study to simulate the membrane potential of twelve distinct types of neurons, each exhibiting a unique spike pattern. The device potential and proteinoid frequency and amplitude were measured for each type of neuron in our study (see supplementary material). The graphical representations in Figures 7-17 elucidate the distinct patterns of the membrane potential, device potential, and proteinoid amplitude across various types of neurons. The data presented in the figures demonstrate that each category of neuron exhibits a discernible spiking pattern, characterised by varying frequencies, amplitudes, shapes, and duration of spikes. The data presented in the figure demonstrates distinct disparities and fluctuations in both the device potential and proteinoid levels across various neuron types. These observations hold potential for characterising the unique spiking behaviour exhibited by each neuron type.

In order to establish a quantitative understanding of the association between the potential of the device and the presence of proteinoids, we conducted an analysis by calculating the cross correlation between these two signals for every distinct neuron type. The cross-correlation is a quantitative metric utilised to assess the degree of similarity between two signals, taking into account the temporal displacement between them. A high cross-correlation value is indicative of a robust linear association between the signals under consideration. Conversely, a low cross-correlation value suggests a feeble or non-existent relationship between the signals. The graphs presented in Figures 7-17 depict the cross correlation analyses conducted between the device potential and the proteinoid potential, for each distinct type of neuron. The presented data illustrates that certain categories of neurons exhibit a notable degree of cross correlation between the mentioned signals, namely tonic bursting and sub-threshold oscillations. Conversely, other neuron types, such as tonic spiking and chattering, demonstrate a comparatively lower level of cross correlation. The findings indicate that certain neuronal sub-types exhibit a higher degree of synchronisation or coherence in their spiking patterns compared to others. Moreover, both the device potential and the presence of proteinoids can serve as reliable indicators of this observed synchronisation or coherence phenomenon.

## 4. Discussion

Proteinoids are synthetic proteins derived from amino acids that can demonstrate spontaneous electrical behaviour. Electrical spikes were observed in an aqueous medium following the synthesis of proteinoids. We demonstrated that L-Glu:L-Phe proteinoids’ electrical activity can be triggered by spiking activity of Izhikevich neurons inputted to the proteinoid ensembles via programmable arbitrary form function generator. Furthermore, the observed asymmetry in the L-Glu:L-Arg proteinoid responses suggests a non-standard interaction between coupled systems. This opens the door to exploring whether proteinoids integrated into bigger networks can display sophisticated collective behaviours.

The findings broaden our comprehension of how amino acid sequence contributes to the generation of spikes in electrical oscillations. Prior research has proposed that distinct amino acid sequences in proteinoids may result in varied electrical oscillation peaks. However, the precise connection between these amino acid variations and peak generation remains incompletely comprehended.

This study demonstrates the essential role of the L-Glu:L-Arg sequence in peak formation, thereby bridging a significant gap in knowledge.

The morphology of proteinoid microspheres and their ensembles, may be influenced by various factors, such as:

**Figure 23:**
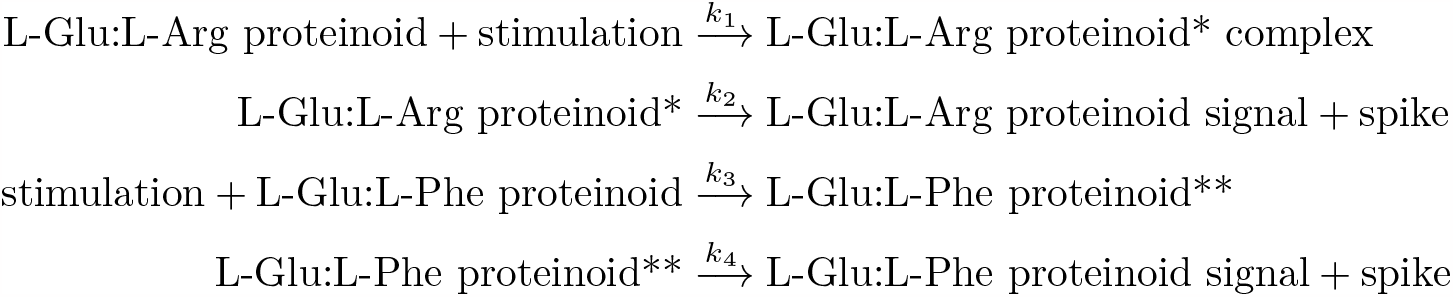
A possible mechanism for the asymmetry in the L-Glu:L-Arg proteinoid responses. Where *k*_1_, *k*_2_, *k*_3_, and *k*_4_ are the rate constants of the reactions.

- The amino acid composition and ratio within the proteinoid mixture [26].
- The pH and temperature of the solution.
- Metal ions, such as calcium, copper, iron, or zinc, can affect the enzyme activities of proteinoids.
- Exposure to electric fields or light can modify the electrical activity of proteinoids [27].

This study utilised the Lempel-Ziv complexity (LZC) to examine the complexity of neuronal dynamics by analysing spike patterns and other variables in the data matrix. The LZC is a metric that quantifies the level of randomness or unpredictability in a sequence. It operates on the principle of identifying and compressing repeated patterns in order to determine the shortest possible description of the sequence. The LZC exhibited variability among various spike patterns, with certain patterns displaying greater complexity and unpredictability. In addition, our findings indicate a lack of significant correlation between the LZC and the variables, with the exception of a modest negative correlation observed between the LZC and device frequency. This implies that the LZC has the ability to capture certain aspects of spiking behaviour that are not directly represented by the variables. Consequently, it can serve as a distinguishing feature for different types of neuronal dynamics. Our findings align with prior research that employed LZC as a metric for assessing the complexity of chaotic systems, fractals, and biomedical signals [24]. Our findings further broaden the utilisation of LZC in the domain of underwater acoustic signal processing, which is a significant and demanding area of study. LZC has potential as a valuable tool for characterising the complexity of underwater acoustic signals and extracting features to enhance the performance of classification and recognition tasks [21].

In the present study, a research study was conducted to analyse the spike train parameters of L-Glu:L-Arg proteinoids under various stimulation modes. These parameters were subsequently compared with the theoretical predictions of the Izhikevich neuron model. In addition to the aforementioned findings, an analysis of the obtained results will be conducted to shed light on the potential ramifications for comprehending the intricate processes of neural coding and computation within proteinoid networks. The coefficient of variation (C) is a statistical measure used to quantify the degree of variability in the inter-spike intervals. A high C value is indicative of irregular or chaotic spiking patterns, whereas a low C value suggests regular or periodic spiking patterns. The data presented in the table 6 indicates that the L-Glu:L-Arg proteinoids exhibit notably elevated C values across various stimulation modes, with the exception of DAP (delayed after-depolarization) and resonator, where C values are comparatively low. The findings indicate that proteinoids composed of L-Glu:L-Arg exhibit a propensity for generating non-uniform spike trains, unless they are subjected to the influence of slow oscillations or resonant frequencies. The aforementioned observation aligns with the well-established Izhikevich neuron model, which posits that distinct configurations of parameter values have the potential to elicit diverse spiking patterns, spanning from regular to chaotic in nature [28]. The measurement known as the fraction of uncorrelated spikes (*F*_*u*_) quantifies the level of independence exhibited by spikes in relation to the stimulation they receive. A positive *F*_*u*_ value signifies the occurrence of spontaneous spike generation, whereas a negative *F*_*u*_ value suggests that the stimulation leads to the suppression of certain spikes. The data presented in the Table 6 indicates that the L-Glu:L-Arg proteinoids exhibit predominantly positive *F*_*u*_ values across various stimulation modes. However, it is noteworthy that the integrator, DAP, inhibition induced bursting, and spike latency stimulation modes are exceptions to this trend, as they display negative *F*_*u*_ values. The findings indicate that the proteinoids composed of L-Glu:L-Arg exhibit a propensity for generating spontaneous spikes, unless subjected to certain conditions such as constant input, slow oscillations, inhibitory input, or delayed input, which appear to modulate their spiking behaviour. This observation aligns with the findings of the Izhikevich neuron model, a computational model that suggests diverse responses to external stimuli can be generated by varying the parameter values. These responses can span from excitatory to inhibitory in nature [29].

**Figure 24:**
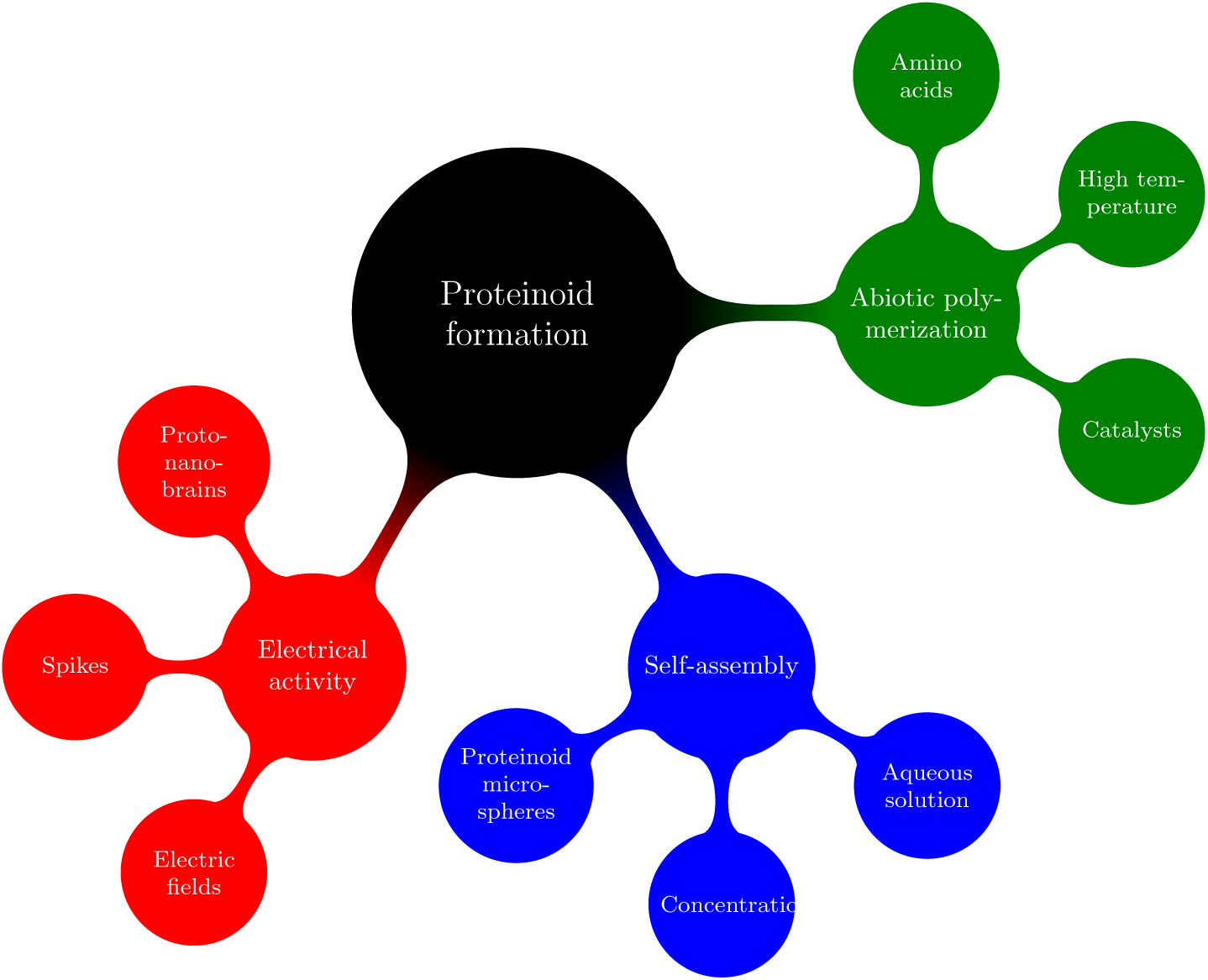
Figure 24: A mind map depicting the mechanisms involved in proteinoid formation and the generation of proto-nano-brains.

The synchrony index (S) quantifies the extent to which the occurrences of spikes align with the timing of the applied stimulation. A higher S value is indicative of a synchronisation between the spikes and the stimulation, whereas a lower S value suggests a desynchronization between the spikes and the stimulation. The data presented in the table 6 indicates that the L-Glu:L-Arg proteinoids exhibit comparatively low S values across the majority of stimulation modes. However, it is noteworthy that the DAP and resonator stimulation modes demonstrate elevated S values for these proteinoids. The findings indicate that proteinoids composed of L-Glu:L-Arg exhibit a propensity for generating desynchronized spike trains, with the exception of instances where they are subjected to slow oscillations or resonant frequencies, which appear to elicit distinct responses. This observation aligns with the findings of the Izhikevich neuron model, a computational model that suggests diverse synchronisation patterns can emerge based on variations in parameter values. These patterns can span from precise phase locking to less precise phase slipping [30]. In conclusion, an extensive analysis has been conducted on the spike train parameters exhibited by L-Glu:L-Arg proteinoids under various stimulation modes. These parameters have been meticulously compared to the theoretical predictions derived from the well-established Izhikevich neuron model. The observed findings indicate that both models exhibit comparable trends and patterns pertaining to spiking variability, spontaneity, and synchrony. The findings of this study indicate that it is plausible to conceptualise the L-Glu:L-Arg proteinoids as a complex system composed of interconnected Izhikevich neurons. Notably, the specific parameter values of these neurons appear to vary depending on the mode of stimulation employed. The proposed framework exhibits potential utility in facilitating the investigation of neural coding and computation within proteinoid networks [31, 32].

## 5. Conclusion

The study on the modulation of electrical activity of ensembles of proteinoid microspheres by simulated Izhikevich neurons has shown that proteinoid microspheres and spikes can be utilised to stimulate inputs and process outputs at different time intervals, thereby providing background for design of neuromorphic devices from thermal proteins, which could be used in unconventional computing, organic electronics and embeddable neuromorphic devices.

## Acknowledgement

The research was supported by EPSRC Grant EP/W010887/1 “Computing with proteinoids”. Authors are grateful to David Paton for helping with SEM imaging and to Neil Phillips for helping with instruments.

